# Determinants of anemia among pregnant women in northern Ghana

**DOI:** 10.1101/708784

**Authors:** Martin N. Adokiya, Richmond Aryeetey, Monica Yost, Andrew D. Jones, Mark L. Wilson

## Abstract

Anemia is a global public health issue affecting half of all pregnant women in developing countries. In 2014, 42% of Ghanaian pregnant women aged 15-49 years were anemic (<11.0g/dl) but information on the determinants of anemia, particularly dietary diversity during the critical third trimester of pregnancy is limited. We assessed the association between determinants and anemia among pregnant women in northern Ghana.

We employed a cross-sectional design involving 624 pregnant women (≥20 weeks of gestation) attending four antenatal care (ANC) health facilities ~25 kilometres north of Tamale, Ghana between July and August 2017. Hemoglobin concentration (measured using Hemocue HB 301) was classified as severe, moderate, or mild. Other data included socio-demographic characteristics, malaria prevention, deworming, and iron/folate tablet use. The FAO Minimum Dietary Diversity (MDD-W) metric was used to categorize women into “inadequate” (MDD-W <5 food groups) and “adequate” (MDD-W ≥5). Logistic regression models were used to determine the association between moderate/severe anemia (Hb<9.0g/dl) and mild anemia (9.0-10.9g/dl), or with ‘no anemia’ (≥11.0g/dl) using STATA 14 software.

Of 624 women sampled, hemoglobin data were available for 601. The mean age was 27.81±0.25 years, gestational age was 31.93±0.13 weeks, ANC attendance was 3.89±0.07; Hb concentration was 9.73g/dl±0.07, and MDD-W index for ten food groups was 5.33±0.04. Anemia (Hb<11.0g/dl) was observed in 74.8% of women (moderate/severe anemia=33.4% and mild anemia=41.4%). Using adjusted logistic regression, women who received deworming medication had lesser odds of being moderate/severe anemic (aOR=0.51, P=0.021). While women who were engaged in other occupation (herdsmen) and no previous parity had higher odds of being moderate/severe anemic (aOR=2.90, P=0.042) and (aOR=2.13, P=0.004) respectively. Moderate/severe anemia was not statistically associated with MDD-W, nor with socioeconomic status/wealth index. Conclusion, anemia in pregnancy was nearly twice that of Ghana as a whole. Deworming medication was found to be protective intervention for anemia during pregnancy.

## INTRODUCTION

Anemia is characterized by low blood hemoglobin (Hb) concentration and constitutes an important public health problem globally. Anemia has both short- and long-term consequences such as preterm, low birth weight, morbidity and mortality ^1,2,3,4^. In 2016, World Health Organization (WHO) estimated that anemia affected 38.2% of pregnant women globally, with the highest prevalence in South-East Asia (48.7%) and Africa (46.3%) ^5,6^. Anemia affects about 1.62 billion people, 56 million of whom are pregnant women ^5^. An estimated 800,000 pregnant women globally have severe anemia (Hb<7.0g/dl). In Ghana, a national Demographic and Health Survey in 2014 determined that 42% of pregnant women were anemic compared to 70% in rural parts of the country ^7^.

Anemia often results from decreased red blood cell production or increased destruction/loss ^3^. Causes include environmental, behavioral, and social factors ^8^ that limit adequate nutrient intake and absorption, or exposure to infectious diseases. In addition, anemia risk is related to household-level factors such as access to water and sanitation, availability of health services, access to diverse food sources, use of insecticide treated nets (ITNs) and knowledge about anemia prevention. Other household- or community-level factors include socioeconomic status, culture, wealth status and education attainment ^5,9,8^.

In developing countries, pregnant women often start gestation with depleted or low body iron stores, making them especially vulnerable to iron deficiency anemia ^6,10,11,12^. Hb concentration declines during pregnancy, partly because of expanded plasma volume compared to red cell mass ^10,13^. This is influenced partly by the iron status of the pregnant woman,^12^ representing a major public health problem in sub-Saharan Africa ^9,10^. Another contributor to anemia is parasitic infections/infestation such as malaria, hookworm and schistosomiasis, especially in areas of Ghana where these infections are endemic ^5^. In addition, chronic infections such as tuberculosis (TB) and human immune-deficiency virus (HIV) increase risk of anemia ^5^. This condition may lead to premature delivery, intrauterine growth retardation, and increased risk of malnutrition, morbidity and mortality for the mother, growing fetus and newborn ^10,11,14^.

Poor maternal diet during preconception and pregnancy is a major contributor to adverse pregnancy outcome such as preterm, low birth weight, still birth and mortality. Diets of pregnant women in developing countries are often limited to a few plant-based foods, with little consumption of micronutrient-dense animal-source foods, or diverse fruits and vegetables ^13,15^. Poor dietary intake and low iron bioavailability are key determinants of low iron reserves and anemia ^16^, particularly with little dietary diversity among poor populations ^17^, who consume mainly carbohydrates with little or no animal products, fruits and vegetables ^10,18^. Little is known about diet and anemia among pregnant women in Ghana. Our study focused on northern Ghanaian women at high-risk of anemia in their third trimester of pregnancy ^7^. We aimed to assess the association between determinants, particularly dietary diversity and moderate/severe anemia, with the hypothesis that, greater dietary diversity would be associated with lower anemia risk.

## Materials and methods

### Study Area/Design

Ghana is a West African nation of 29 million people who are mostly concentrated in the southern and coastal regions. Economically, Ghana ranks in the top third of African nations (GDP/capita = US$ 4,600), with considerable geographic variation in wealth. Our study was conducted in Northern Region, a poorer, predominantly agricultural area, made up of 28 Districts and Municipalities. Four government antenatal care (ANC) health facilities in Savelugu Municipality, located ~ 25 kilometres north of Tamale, served as the source of the study women. We employed a cross-sectional design involving pregnant women (≥20 weeks of gestation) seeking ANC, using our own questionnaire and ANC medical record data to identify risk factors of anemia. The study was undertaken during July-August 2017 (rainy season), using a questionnaire that had been pre-tested at a nearby health center.

### Study population and protection of human subjects

Data were collected from pregnant women attending health facilities in the North, South, East and West quadrants of the Municipality. One health facility (HF) was randomly selected in the South (Janjori Kukuo Health Centre) to pre-test the tools/questionnaire. Then, a HF was randomly selected from each of the remaining three quadrants: Moglaa Health Centre (West), Savelugu Reproductive and Child Health (East) and Pong Tamale Health Centre (North). The fourth HF sampled was Savelugu Hospital, a major referral and health-seeking hospital centrally located in the Municipality that serves many ANC-seeking women. During the ANC days of the four HFs, pregnant women were recruited following informed consent. Women attending the ANC were informed of the study in the native language, Dagbani, by a member of the study team. Women were eligible to participate if they were at the HF to receive ANC, pregnant with a gestational age of at least 20 weeks, 18 or more years of age, and had not been diagnosed with sickle cell anemia. The maternal records of interested pregnant women were examined for compliance to these inclusion criteria. For eligible women, a sticker with a unique identification number was placed on their maternal record book and they were invited to stay for an interview. The women were then seen one-on-one with a trained interviewer who explained the details of the study, including risks and benefits. Women who agreed to participate gave their consent via thumb print, and were then provided with a signed copy of their consent form in English, as the native language is not commonly written, and many people are illiterate. The study protocol was approved before its implementation by the Ghana Health Service Ethical Review Committee (GHSERC/12/05/17) and the University of Michigan Institutional Review Board for Health Sciences and Behavioral Sciences (HUM00128583). Additionally, official approval letters were obtained from the Regional Director and District Director of Ghana Health Service in Northern Region and Savelugu Municipality, respectively, as well as heads of the four HFs.

### Data sources and derived variables

Data were gathered from oral interviews using a pretested questionnaire, and from the ANC record of each woman. The outcome of interest was the pregnant woman’s anemia status. ANC is mostly free to pregnant women in Ghana, and provides various interventions and preventative care to combat anemia and infections ^7^. Public health service procedures ensure all pregnant women have their hemoglobin tested to monitor anemia during each ANC visit, and receive iron supplementation, intermittent preventive treatment of malaria during pregnancy (IPTp), deworming medication, insecticide treated bed nets (ITN), and education about diet and malaria prevention ^7^. While these services are, in theory, widely available to all pregnant women, there are many factors influencing whether a woman actually receives this care. Even though the ANC services are free, the transportation to the clinic is not. Some women have to walk for many miles to reach the nearest ANC center. Rural health centers are often poorly staffed and poorly supplied with materials and medications needed for the services. Additionally, attending ANC may mean the loss of a day’s income, which may not be possible for lower-income women. Despite these challenges, a recent study reported that 97% of pregnant Ghanaian women attended at least one ANC visit in 2014, and 87% had attended four or more visits ^7^. In the Northern Region where our study occurred, ANC attendance is likely to be lower due to lower income and fewer health centers. Even with high ANC attendance, anemia remains a major problem for adult women in many regions of Ghana, particularly in areas like our study sites where poverty levels are high, and access to health centers is hindered ^7^.

The interviews were conducted by trained local research assistants with health education backgrounds (e.g. Bachelor of Science degree in Nursing or Community Nutrition). Each hired research assistant was fluent in the Dagbani and English languages, as well as other local native Ghanaian languages. The interviews were conducted in a private room or area of the HF to protect women’s privacy. Individual women were asked questions concerning their demographic situation, as well as characteristics of their housing, water, toilets, and household assets. In addition, they were asked about the ANC services they had received, and were administered a 24-hour dietary recall survey, collecting information on all food items and beverages consumed in the previous day. Interviewers asked individual women to recall all foods they had consumed in the previous 24-hour, and after responding were probed to ensure that no meal or snack was left out (breakfast, snack before lunch, lunch, snack after lunch, dinner and snack before going to bed). The foods were then categorized into ten (10) food groups ^19^. Information on Hb concentration, and gestation, were extracted from the maternal health records that are kept for each pregnant woman.

Hb concentration determination differed between the HFs. The Savelugu hospital used a spectrophotometer operated by trained laboratory scientist/technicians. The three health centers used a Hemocue HB 301 operated by HF workers. A blood sample for Hb concentration was taken on the day of the interview in most cases, but for very few participants this occurred one week after the interview.

Our study used results from tests that are routinely conducted at ANC visits, as recommended by the Government of Ghana. All tests were performed by HF staff. Occasionally, tests were not performed on the day of interview due to shortages of supplies. However, tests for these women were completed in the following weeks if they returned to the same HF. Overall, 27 women did not return for testing. After completing the interview and data extraction, participants were given GHS10.00 (US$ 2.27) to compensate them for transportation back to their homes.

### Statistical analysis

Data were cleaned and analyzed using STATA 14 software. Frequency tables were generated to describe the distribution of anemia status and all hypothesized explanatory variables. Logistic regression was performed to determine the unadjusted associations between anemia status and each independent explanatory variable. Adjusted logistic regression analyses were performed to evaluate relationships among independent factors and anemia status. As with the Demographic and Health Survey’s (DHS) classification scheme, we defined anemia as severe (Hb<7.0g/dl), moderate (Hb =7.0-8.9g/dl), or mild (Hb=9.0-10.9g/dl), with no anemia being Hb≥11.0g/dl. This cut-off was selected in accordance with the Demographic and Health Survey for Ghana and previous studies of anemia in pregnant women ^3,7,10^. For our study, severe and moderate anemia were combined (moderate/severe) as Hb <9.0g/dl, and analyzed in comparison with mild/no anemia. Anemia prevalence was analyzed for the entire study population, as well as for each HF, using logistic regression tests to evaluate risk factor associations.

Dietary diversity was calculated using the reported number of different food groups consumed by each woman in the previous 24 hours. Food was reclassified into ten distinct food groups: (1) grains, roots and tubers, (2) Pulses-beans, peas and lentils, (3) Nuts and seeds, (4) Dairy, (5) Meat, poultry and fish, (6) Eggs, (7) Dark green leafy vegetables, (8) Vitamin A-rich fruits and vegetables, (9) Other vegetables and (10) Other fruits. These mutually exclusive food groups are those that comprise the FAO Minimum Dietary Diversity for Women (MDD-W) indicator ^19^. The FAO MDD-W metric was used to categorize participants into “inadequate” (MDD-W <5 food groups) and “adequate” (MDD-W ≥5). A wealth index was calculated using Principal Component Analysis (PCA) of household assets, housing conditions, water facilities and toilet facilities. Other risk factors that were analyzed included maternal education, religion, ethnicity, and occupation as well as socio-demographic characteristics, ANC attendance, parity, ITN ownership and ITN utilization. The relation between each individual risk factor and moderate/severe anemia was determined using bivariate logistic regression to obtain the odds ratios (ORs), confidence intervals (CIs) and probability values (p-value) for the whole sample. Separate adjusted logistic regression models were constructed for the whole sample using ten (10) selected risk factors with a significance test level of alpha = 0.05. Odds ratios were used to interpret the associations of risk factors with moderate/severe or mild anemia/no anemia.

## RESULTS

### Study population characteristics

In total, 624 pregnant women at ≥20 weeks of gestation were enrolled, but analysis was limited to 601 participants (96.3%) for whom Hb measurements were available (**Table 1)**. The mean age was 27.81 (±0.25) years, with gestation age =31.93 (±0.13), ANC= 3.89 (±0.07), Hb =9.73 (±0.07) and MDD-W = 5.33 (±0.04). More than half (58%) of the participants who received ANC services at the Savelugu Hospital and 53% were between 20 and 29 years old. Nearly all women (96%) were from the Dagomba ethnic group. Three-quarters (75%) had no formal education, and nearly all (97%) belonged to the Islamic religion, with 98% being married (monogamous or polygamous marriages). About 69% of the participants were either farmers (39%) or petty traders (30%), while very few (2%) were salaried workers. Most women (91%) were between 28 and 36 weeks of gestation when interviewed, while very few (2%) were between 20 and 27 weeks of gestation. Approximately 92% of the participants had access to improved water sources and about one-quarter (24%) reported using improved toilet facilities.

**Table 1:** Socio-demographic factors and dietary diversity by anemia status among pregnant women in northern Ghana (n=601)

| Variables | Total Sample<br>(N=601) | Moderate/severe<br>anemia<br>(<9.0g/dl) | Mild anemia<br>(9.0-10.9g/dl) | No anemia<br>(≥11.0g/dl) |
| --- | --- | --- | --- | --- |
|  | Frequency (%) | Frequency (%) | Frequency (%) | Frequency (%) |
| **Name of Health Facility** |  |  |  |  |
| Savelugu hospital | 349(58.1) | 157(45.0) | 108(31.0) | 84(24.0) |
| Moglaa HC | 93(15.5) | 14(15.0) | 54(58.1) | 25(26.9) |
| Pong Tamale HC | 77(12.8) | 18(23.4) | 44(57.1) | 15(19.5) |
| Savelugu RCH | 82(13.6) | 12(14.6) | 43(52.4) | 27(33.0) |
| **Age (in years)** |  |  |  |  |
| <20 | 34(5.7) | 14 (41.2) | 15(44.1) | 5(14.7) |
| 20-29 | 319(53.1) | 112(35.1) | 125(39.2) | 82(25.7) |
| 30-39 | 226(37.6) | 73(32.3) | 95(42.0) | 58(25.7) |
| ≥40 | 22(3.6) | 2(9.1) | 14(63.6) | 6 (27.3) |
| **Ethnicity** |  |  |  |  |
| Dagomba | 576(96.2) | 195(33.8) | 236 (41.0) | 145(25.2) |
| Fulani | 8(1.3) | 2(25.0) | 5(62.5) | 1(12.5) |
| Others (Akan, Ewe etc.) | 15(2.5) | 3(20.0) | 7(46.7) | 5(33.3) |
| **Education status** |  |  |  |  |
| None | 450(74.9) | 139(30.9) | 204(45.3) | 107(23.8) |
| Primary | 46(7.7) | 19(41.3) | 16(34.8) | 11(23.9) |
| JSS/Middle School | 53(8.8) | 22(41.5) | 15(28.3) | 16(30.2) |
| Senior Secondary | 38(6.3) | 17(44.7) | 11(29.0) | 10(26.3) |
| Tertiary | 14(2.3) | 4(28.6) | 3(21.4) | 7(50.0) |
| **Religious denomination** |  |  |  |  |
| Islam | 582(97.3) | 196(33.7) | 238(40.9) | 148(25.4) |
| Christianity | 13(2.2) | 4(30.8) | 7(53.8) | 2(15.4) |
| Traditionalist | 3(0.50) | 0(0.0) | 2(66.7) | 1(33.3) |
| **Marital status** |  |  |  |  |
| Single | 12(2.0) | 5(41.7) | 7(58.3) | 0(0.0) |
| Married-monogamous | 329(54.7) | 103(31.3) | 139(42.3) | 87(26.4) |
| Married-polygamous | 260(43.3) | 93(35.8) | 103(39.6) | 64(24.6) |
| **Occupation status** |  |  |  |  |
| Farmer | 233(38.8) | 73(31.3) | 108(46.4) | 52(22.3) |
| Petty trader | 181(30.1) | 54(29.9) | 73(40.3) | 54(29.8) |
| Salaried worker | 14(2.3) | 4(28.6) | 4(28.6) | 6(42.8) |
| Artisan/laborer | 79(13.1) | 27(34.2) | 34(43.0) | 18(22.8) |
| No occupation | 75(12.5) | 31(41.3) | 27(36.0) | 17(22.7) |
| Others | 19(3.2) | 12(63.2) | 3(15.8) | 4(21.0) |
| **Wealth index** |  |  |  |  |
| Poorest | 118(19.6) | 34(28.8) | 53(44.9) | 31(26.3) |
| Poor | 119(19.8) | 41(37.5) | 50(42.0) | 28(23.5) |
| Medium | 120(20.0) | 45(37.5) | 51(42.5) | 24(20.0) |
| Wealthy | 122(20.3) | 38(31.1) | 45(36.9) | 39(32.0) |
| Wealthiest | 122(20.3) | 43(35.2) | 50(41.0) | 29(23.8) |
| **Gestational age (in weeks)** |  |  |  |  |
| 20-27 | 11(1.8) | 4(36.4) | 4(36.4) | 3(27.2) |
| 28-36 | 547(91.0) | 182(33.2) | 229(41.9) | 136(24.9) |
| ≥37 | 43(7.2) | 15(34.9) | 16(37.2) | 12(27.9) |
| **Drinking water source** |  |  |  |  |
| Non-improved | 48(8.0) | 12(25.0) | 25(52.1) | 11(22.9) |
| Improved | 553(92.0) | 189(34.2) | 224(40.5) | 140(25.3) |
| **Toilet facility** |  |  |  |  |
| Non-improved | 457(76.0) | 155(33.9) | 185(40.5) | 117(25.6) |
| Improved | 144(24.0) | 46(31.94) | 64(44.44) | 34(23.61) |
| **Grains/roots/tubers/plantain** |  |  |  |  |
| No | 1(0.2) |  |  |  |
| Yes | 600(99.8) |  |  |  |
| **Pulses** |  |  |  |  |
| No | 452(75.3) | 139(30.8) | 194(42.9) | 119(26.3) |
| Yes | 148(24.7) | 61(41.2) | 55(37.2) | 32(21.6) |
| **Nuts/seeds** |  |  |  |  |
| No | 133(22.2) | 48(36.1) | 53(39.8) | 32(24.1) |
| Yes | 467(77.8) | 153(32.8) | 195(41.7) | 119(25.5) |
| **Dairy** |  |  |  |  |
| No | 477(79.4) | 158(33.1) | 197(41.3) | 122(25.6) |
| Yes | 124(20.6) | 43(34.7) | 52(41.9) | 29(23.4) |
| **Meat/poultry/fish(amaani)** |  |  |  |  |
| No | 7(1.2) | 2(28.6) | 3(42.8) | 2(28.6) |
| Yes | 594(98.8) | 199(33.5) | 246(41.4) | 149(25.1) |
| **Eggs** |  |  |  |  |
| No | 555(92.5) | 185(33.3) | 233(42.0) | 137(24.7) |
| Yes | 45(7.5) | 16(35.6) | 16(35.6) | 13(28.8) |
| **Dark green leafy vegetables** |  |  |  |  |
| No | 40(6.7) | 10(25.0) | 22(55.0) | 8(20.0) |
| Yes | 561(93.3) | 191(34.0) | 227(40.5) | 143(25.5) |
| **Vitamin A-rich fruits & vegetables** |  |  |  |  |
| No | 555(92.3) | 192(34.6) | 223(40.2) | 140(25.2) |
| Yes | 46(7.7) | 9(19.6) | 26(56.5) | 11(23.9) |
| **Other vegetables** |  |  |  |  |
| No | 11(1.8) | 3(27.3) | 6(54.5) | 2(18.2) |
| Yes | 590(98.2) | 198(33.6) | 243(41.2) | 149(25.2) |
| **Other fruits** |  |  |  |  |
| No | 570(95.0) | 191(33.5) | 234(41.1) | 145(25.4) |
| Yes | 30(5.0) | 9(30.0) | 15(50.0) | 6(20.0) |
| **Minimum dietary diversity score** |  |  |  |  |
| <5 | 91(15.1) | 27(29.7) | 40(43.9) | 24(26.4) |
| ≥5 | 510(84.9) | 174(34.1) | 209(41.0) | 127(24.9) |
| **Household livestock** |  |  |  |  |
| No | 116(19.3) | 36(31.0) | 45(38.8) | 35(30.2) |
| Yes | 485(80.7) | 165 (34.0) | 204(42.1) | 116(23.9) |
| **Household ITN** |  |  |  |  |
| No | 55(9.2) | 27(49.1) | 17(30.9) | 11(20.0) |
| Yes | 546(90.8) | 174(31.9) | 232(42.5) | 140(25.6) |
| **Slept under ITN previous night** |  |  |  |  |
| No | 104(17.3) | 46(44.2) | 35(33.7) | 23(22.1) |
| Yes | 497(82.7) | 155(31.2) | 214(43.1) | 128(25.7) |
| **No. of previous deliveries** |  |  |  |  |
| 0 | 142(23.6) | 68(47.9) | 48(33.8) | 26(18.3) |
| 1 | 122(20.3) | 40(32.8) | 54(44.3) | 28(22.9) |
| 2-3 | 214(35.6) | 59(27.6) | 91(42.5) | 64(29.9) |
| ≥4 | 123(20.5) | 34(27.7) | 56(45.5) | 33(26.8) |
| **Total ANC visits** |  |  |  |  |
| < 4 | 286(47.6) | 97(33.9) | 123(43.0) | 66(23.1) |
| 4-7 | 286(47.6) | 96(33.6) | 111(38.8) | 79(27.6) |
| ≥8 | 29(4.8) | 8(27.6) | 15(51.7) | 6(20.7) |
| **IPT received** |  |  |  |  |
| No | 99(16.5) | 41(41.4) | 38(38.4) | 20(20.2) |
| Yes | 502(83.5) | 160(31.9) | 211(42.0) | 131(26.1) |
| **Deworming medication** |  |  |  |  |
| No | 515(85.7) | 183(35.5) | 198(38.5) | 134(26.0) |
| Yes | 86(14.3) | 18(20.9) | 51(59.3) | 17(19.8) |
| **Received iron tablets** |  |  |  |  |
| No | 3(0.5) | 1(33.3) | 1(33.3) | 1(33.3) |
| Yes | 598(99.5) | 200(33.4) | 248(41.5) | 150(25.1) |
| **Received folic acid** |  |  |  |  |
| No | 3(0.5) | 1(33.3) | 1(33.3) | 1(33.3) |
| Yes | 598(99.5) | 200(33.4) | 248(41.5) | 150(25.1) |
| **Prevalence of anemia** |  |  |  |  |
| Severe-moderate (<9.0g/dl) | 201(33.5) |  |  |  |
| Mild (9.0-10.9g/dl) | 249(41.4) |  |  |  |
| No anemia | 151(25.1) |  |  |  |
| **Prevalence of anemia types** |  |  |  |  |
| Severe (<7.0g/dl) | 25(4.2) |  |  |  |
| Moderate (7.0-8.9g/dl) | 175(29.2) |  |  |  |
| Mild (9.0-10.9g/dl) | 249(41.5) |  |  |  |
| No anemia (≥11.0g/dl) | 151(25.1) |  |  |  |
| **Means** | **Mean (CI)** | **Standard Error** |  |  |
| Age (in years) | 27.81(27.33, 28.29) | 0.246 |  |  |
| Gestational age (in weeks) | 31.93(31.68, 32.18) | 0.128 |  |  |
| Number of ANC visits | 3.89(3.76, 4.03) | 0.070 |  |  |
| Hemoglobin level | 9.73(9.59, 9.88) | 0.074 |  |  |
| Minimum Dietary diversity | 5.33(5.25, 5.42) | 0.042 |  |  |

About 85% of the participants achieved the MDD-W indicator, that is, consumed five or more food groups in the previous 24 hours. In addition, 91% of the participants reported household ownership of at least one ITN, with 83% having slept under an ITN on the night preceding the interview. About three-quarters (76%) of the participants had previously delivered live babies. While 52% had attended ≥4 ANC visits and 84% received at least one IPTp treatment, only 14% received any deworming medication. In addition, 99% of the participants reported they had received iron and folic acid tablets during ANC visits.

### Prevalence of moderate/severe anemia and risk factors

Three-quarters (75%) of the pregnant women in the study were anemic with 41% experiencing mild anemia and 33% experiencing moderate/severe anemia. Forty-one percent of participants who were <20 years of age had moderate/severe anemia. Anemia prevalence declined with increasing age. Moderate/severe anemia was least prevalent among salaried workers (29%) and petty traders (30%), but highest among unemployed (41%) and others, such as herdsmen (63%).

Nearly half (49%) of the participants without household ownership of ITNs were moderate/severe anemic compared to 32% among those with ITNs. Those who did not sleep under ITN on the previous night preceding the survey were more likely to experience moderate/severe anemia (44%) compared to those who slept under an ITN (31%). In addition, 41% of the participants who did not receive Intermittent Preventive Treatment (IPT) had moderate/severe anemia compared to 32% in those who did receive IPT.

Women with no previous delivery had the greater risk of moderate/severe anemia (48%) compared to women with at least one previous deliveries. More than a third of pregnant women who did not receive deworming medication had moderate/severe anemia compared to about one-fifth among those who did take deworming drugs. Similarly, about 33% of the participants who received iron tablets or folic acid tablets did not have moderate/severe anemia (**Table 1**).

Participants who consumed pulses (41%), dairy (35%), meat, poultry and fish (34%), eggs (36%), dark green leafy vegetables (34%), and other vegetables (34%) had greater risk of moderate/severe anemia than those who did not. However, participants who consumed nuts/seeds (33%), vitamin A-rich fruits and vegetables (20%) and other fruits (30%) were less likely to have moderate/severe anemia compared to those who did not consume.

### Analysis of risk factors and any anemia or moderate/severe anemia

When all three levels of anemia were combined (<11.0g/dl) and compared with no anemia using bivariate (unadjusted) logistic regression (**Table 2**), anemia risk was significantly lower with tertiary education (**OR=0.31**, 95%CI:0.11, 0.91), and higher with medium wealth (**OR=1.88**, 95%CI:1.04, 3.38) and greater parity (**OR=1.90,**95%CI:1.14, 3.19). However, when comparison involved women classified as moderate/severe anemic (<9.0g/dl) versus combined mild/no anemia, nine (9) explanatory variables were significantly associated. Women seeking ANC services at Savelugu Hospital were more likely to have moderate/severe anemia than those at the other three HFs, including Moglaa HC (**OR=0.22**, 95%CI:0.12, 0.40), Pong Tamale HC (**OR=0.37**, 95%CI:0.21, 0.66) and Savelugu RCH (**OR=0.21**, 95%CI:0.11, 0.40). Moderate/severe anemia was less frequent among women who were older (**OR=0.18,** 95%CI:0.04, 0.81). Participants who consumed vitamin A-rich fruits and vegetables were less likely to have moderate/severe anemia than those who did not consume (**OR=0.46,** 95%CI:0.22, 0.97). Similarly, women who received deworming medication were at lower risk to have moderate/severe anemia than those who did not receive (**OR=0.48**, 95%CI:0.28, 0.83). Interestingly, moderate/severe anemia was more likely among women who were engaged in other occupations (e.g. herdsmen) (**OR=3.76**, 95%CI:1.42, 9.94), or who consumed pulses (**OR=1.58,** 95%CI:1.06, 2.32).

**Table 2:** Bivariate odd ratios for anemia status among pregnant women in Northern Ghana (n=601)

| | | Anemia<br>($<11.0\text{g/dl}$) | No<br>anemia<br>($\geq 11.0\text{g/dl}$) | Anemia ($<11.0\text{g/dl}$) versus<br>no anemia ($\geq 11.0\text{g/dl}$) | | Moderate/<br>severe<br>anemia<br>($<9.0\text{g/dl}$) | Mild/no<br>anemia<br>($\geq 9.0\text{g/dl}$) | Moderate severe anemia<br>($<9.0\text{g/dl}$) versus mild/no<br>anemia ($\geq 9.0\text{g/dl}$) | |
| --- | --- | --- | --- | --- | --- | --- | --- | --- | --- |
|  | Frequency | % | % | unadjusted OR(CI) | P-value | % | % | unadjusted<br>OR(CI) | P-value |
| **Health Facility** |  |  |  |  |  |  |  |  |  |
| Savelugu hospital | 349 | 75.9 | 24.1 | Ref |  | 45.0 | 55.0 | Ref |  |
| Moglaa HC | 93 | 73.1 | 26.9 | 0.86(0.51, 1.45) | 0.576 | 15.1 | 84.9 | 0.22(0.12, 0.40) | **<0.001** |
| Pong Tamale HC | 77 | 80.5 | 19.5 | 1.31(0.71, 2.42) | 0.389 | 23.4 | 76.6 | 0.37(0.21, 0.66) | **0.001** |
| Savelugu RCH | 82 | 67.1 | 32.9 | 0.65(0.38, 1.09) | 0.100 | 14.6 | 85.4 | 0.21(0.11, 0.40) | **<0.001** |
| **Age (in years)** |  |  |  |  |  |  |  |  |  |
| $<20$ | 34 | 85.3 | 14.7 | 2.01(0.75, 5.36) | 0.164 | 41.2 | 58.8 | 1.29 (0.63, 2.66) | 0.484 |
| 20-29 | 319 | 74.3 | 25.7 | Ref |  | 35.1 | 64.9 | Ref |  |
| 30-39 | 226 | 74.3 | 25.7 | 1.00(0.68, 1.48) | 0.991 | 32.3 | 67.7 | 0.88(0.61, 1.27) | 0.495 |
| $\geq 40$ | 22 | 72.7 | 27.3 | 0.92(0.35, 2.44) | 0.871 | 9.1 | 90.9 | 0.18(0.04, 0.81) | **0.025** |
| **Ethnicity** |  |  |  |  |  |  |  |  |  |
| Dagomba | 576 | 74.8 | 25.2 | Ref |  | 33.8 | 66.2 | Ref |  |
| Fulani | 8 | 87.5 | 12.5 | 2.35(0.28, 19.30) | 0.425 | 25.0 | 75.0 | 0.65(0.13, 3.26) | 0.602 |
| Others (e.g. Akan, Ewe<br>etc.) | 15 | 66.7 | 33.3 | 0.67(0.23, 2.00) | 0.476 | 20.0 | 80.0 | 0.49(0.14, 1.75) | 0.271 |
| **Education status** |  |  |  |  |  |  |  |  |  |
| None | 450 | 76.2 | 23.8 | Ref |  | 30.9 | 69.1 | Ref |  |
| Primary | 46 | 76.1 | 23.9 | 0.99(0.49, 2.02) | 0.984 | 41.3 | 58.7 | 1.57(0.85, 2.93) | 0.151 |
| JSS/Middle School | 53 | 69.8 | 30.2 | 0.72(0.39, 1.35) | 0.306 | 41.5 | 58.5 | 1.59(0.89, 2.84) | 0.119 |
| Senior Secondary | 38 | 73.7 | 26.3 | 0.87(0.41, 1.86) | 0.725 | 44.7 | 55.3 | 1.81(0.93, 3.54) | 0.082 |
| Tertiary | 14 | 50.0 | 50.0 | 0.31(0.11, 0.91) | **0.033** | 28.6 | 71.4 | 0.89(0.28, 2.90) | 0.853 |
| **Occupation status** |  |  |  |  |  |  |  |  |  |
| Farmer | 233 | 77.7 | 22.3 | Ref |  | 31.3 | 68.7 | Ref |  |
| Petty trader | 181 | 70.2 | 29.8 | 0.68 (0.43, 1.05) | 0.083 | 29.8 | 70.2 | 0.93(0.61, 1.42) | 0.743 |
| Salaried worker | 14 | 57.1 | 42.9 | 0.38(0.13, 1.15) | 0.088 | 28.6 | 71.4 | 0.88(0.27, 2.89) | 0.829 |
| Artisan/laborer | 79 | 77.2 | 22.8 | 0.97(0.53, 1.79) | 0.931 | 34.2 | 65.8 | 1.14(0.66, 1.96) | 0.640 |
| No occupation | 75 | 77.3 | 22.7 | 0.98(0.53, 1.83) | 0.950 | 41.3 | 58.7 | 1.54(0.90, 2.64) | 0.112 |
| Others | 19 | 78.9 | 21.1 | 1.08(0.34, 3.39) | 0.899 | 63.2 | 36.8 | 3.76(1.42, 9.94) | **0.008** |
| **Wealth index** |  |  |  |  |  |  |  |  |  |
| Poorest | 118 | 73.7 | 26.3 | 1.32(0.75, 2.31) | 0.332 | 28.8 | 71.2 | 0.89(0.51, 1.56) | 0.693 |
| Poor | 119 | 76.5 | 23.5 | 1.53(0.86, 2.70) | 0.145 | 34.5 | 65.5 | 1.16(0.68, 1.99) | 0.585 |
| Medium | 120 | 80.0 | 20.0 | 1.88(1.04, 3.38) | **0.035** | 37.5 | 62.5 | 1.33(0.78, 2.26) | 0.298 |
| Wealthy | 122 | 68.0 | 32.0 | Ref |  | 31.2 | 68.8 | Ref |  |
| Wealthiest | 122 | 76.2 | 23.8 | 1.51(0.86, 2.65) | 0.154 | 35.3 | 64.7 | 1.20(0.71, 2.05) | 0.497 |
| **Gestational age (in weeks)** |  |  |  |  |  |  |  |  |  |
| 20-27 | 11 | 72.7 | 27.3 | 0.88(0.23,3.37) | 0.855 | 36.4 | 63.6 | 1.15(0.33, 3.97) | 0.830 |
| 28-36 | 547 | 75.1 | 24.9 | Ref |  | 33.3 | 66.7 | Ref |  |
| ≥37 | 43 | 72.1 | 27.9 | 0.85(0.43, 1.71) | 0.658 | 34.9 | 65.1 | 1.07(0.56, 2.06) | 0.829 |
| **Drinking water source** |  |  |  |  |  |  |  |  |  |
| Non-improved | 48 | 77.1 | 22.9 | 1.14(0.57, 2.30) | 0.713 | 25.0 | 75.0 | 0.64(0.33, 1.26) | 0.199 |
| Improved | 553 | 74.7 | 25.3 | Ref |  | 34.2 | 65.8 | Ref |  |
| **Toilet facility** |  |  |  |  |  |  |  |  |  |
| Non-improved | 457 | 74.4 | 25.6 | Ref |  | 33.9 | 66.1 | Ref |  |
| Improved | 144 | 76.4 | 23.6 | 1.11(0.72, 1.73) | 0.631 | 31.9 | 68.1 | 0.91(0.61, 1.36) | 0.662 |
| **Pulses** |  |  |  |  |  |  |  |  |  |
| No | 452 | 73.7 | 26.3 | Ref |  | 30.7 | 69.3 | Ref |  |
| Yes | 148 | 78.4 | 21.6 | 1.30 (0.83, 2.02) | 0.253 | 41.2 | 58.8 | 1.58(1.06, 2.32) | **0.020** |
| **Nuts/seeds** |  |  |  |  |  |  |  |  |  |
| No | 133 | 75.9 | 24.1 | 1.08(0.69, 1.69) | 0.739 | 36.1 | 63.9 | 1.16(0.77, 1.73) | 0.473 |
| Yes | 467 | 74.5 | 25.5 | Ref |  | 32.8 | 67.2 | Ref |  |
| **Dairy** |  |  |  |  |  |  |  |  |  |
| No | 477 | 74.4 | 25.6 | Ref |  | 33.1 | 66.9 | Ref |  |
| Yes | 124 | 76.6 | 23.4 | 1.13 (0.71, 1.79) | 0.617 | 34.7 | 65.3 | 1.07(0.71, 1.62) | 0.744 |
| **Meat/poultry/fish(amaani)** |  |  |  |  |  |  |  |  |  |
| No | 7 | 71.4 | 28.6 | 0.84(0.16, 4.36) | 0.833 | 28.6 | 71.4 | 0.79(0.15, 4.13) | 0.784 |
| Yes | 594 | 74.9 | 25.1 | Ref |  | 33.5 | 66.5 | Ref |  |
| **Eggs** |  |  |  |  |  |  |  |  |  |
| No | 555 | 75.3 | 24.7 | Ref |  | 33.3 | 66.7 | Ref |  |
| Yes | 45 | 71.1 | 28.9 | 0.81(0.41, 1.58) | 0.532 | 35.6 | 64.4 | 1.10(0.58, 2.08) | 0.761 |
| **Dark green leafy vegetables** |  |  |  |  |  |  |  |  |  |
| No | 40 | 80.0 | 20.0 | 1.37(0.62, 3.04) | 0.441 | 25.0 | 75.0 | 0.65(0.31, 1.35) | 0.245 |
| Yes | 561 | 74.5 | 25.5 | Ref |  | 34.1 | 65.9 | Ref |  |
| **Vitamin A-rich fruits & vegetables** |  |  |  |  |  |  |  |  |  |
| No | 555 | 74.8 | 25.3 | Ref |  | 34.6 | 65.4 | Ref |  |
| Yes | 46 | 76.1 | 23.9 | 1.07(0.53, 2.17) | 0.844 | 19.6 | 80.4 | 0.46(0.22, 0.97) | **0.042** |
| **Other vegetables** |  |  |  |  |  |  |  |  |  |
| No | 11 | 81.8 | 18.2 | 1.52(0.32, 7.12) | 0.595 | 27.3 | 72.7 | 0.74(0.19, 2.83) | 0.663 |
| Yes | 590 | 74.7 | 25.3 | Ref |  | 33.6 | 66.4 | Ref |  |
| **Other fruits** |  |  |  |  |  |  |  |  |  |
| No | 570 | 74.6 | 25.4 | Ref |  | 33.5 | 66.5 | Ref |  |
| Yes | 30 | 80.0 | 20.0 | 1.36(0.54, 3.40) | 0.505 | 30.0 | 70.0 | 0.85(0.38, 1.89) | 0.691 |
| **Dietary diversity** |  |  |  |  |  |  |  |  |  |
| <5 | 91 | 73.6 | 26.4 | 0.93(0.56, 1.54) | 0.766 | 29.7 | 70.3 | 0.81(0.50, 1.32) | 0.408 |
| ≥5 | 510 | 75.1 | 24.9 | Ref |  | 34.1 | 65.9 | Ref |  |
| **Household livestock** |  |  |  |  |  |  |  |  |  |
| No | 116 | 69.8 | 30.2 | 0.73(0.46, 1.14) | 0.164 | 31.0 | 69.0 | 0.87(0.56, 1.35) | 0.540 |
| Yes | 485 | 76.1 | 23.9 | Ref |  | 34.0 | 66.0 | Ref |  |
| **Household ITN** |  |  |  |  |  |  |  |  |  |
| No | 55 | 80.0 | 20.0 | 1.38(0.69, 2.74) | 0.360 | 49.1 | 50.9 | 2.06(1.18, 3.60) | **0.011** |
| Yes | 546 | 74.4 | 25.6 | Ref |  | 31.9 | 68.1 | Ref |  |
| **Slept under ITN previous night** |  |  |  |  |  |  |  |  |  |
| No | 104 | 77.9 | 22.1 | 1.22(0.74, 2.02) | 0.437 | 44.2 | 55.8 | 1.75(1.14, 2.69) | **0.011** |
| Yes | 497 | 74.3 | 25.7 | Ref |  | 31.2 | 68.8 | Ref |  |
| **No. of previous deliveries** |  |  |  |  |  |  |  |  |  |
| 0 | 142 | 81.7 | 18.3 | 1.90(1.14, 3.19) | **0.015** | 47.9 | 52.1 | 2.41(1.55, 3.77) | **<0.001** |
| 1 | 122 | 77.1 | 22.9 | 1.43(0.86, 2.39) | 0.170 | 32.8 | 67.2 | 1.28(0.79, 2.08) | 0.314 |
| 2-3 | 214 | 70.1 | 29.9 | Ref |  | 27.6 | 72.4 | Ref |  |
| ≥4 | 123 | 73.2 | 26.8 | 1.16(0.71, 1.91) | 0.548 | 27.64 | 72.36 | 1.00(0.61, 1.65) | 0.989 |
| **Total ANC visits** |  |  |  |  |  |  |  |  |  |
| < 4 | 286 | 76.9 | 23.1 | Ref |  | 33.9 | 66.1 | Ref |  |
| 4-7 | 286 | 72.4 | 27.6 | 0.79(0.54, 1.15) | 0.212 | 33.6 | 66.4 | 0.98(0.70, 1.39) | 0.930 |
| ≥8 | 29 | 79.3 | 20.7 | 1.15(0.45, 2.94) | 0.771 | 27.6 | 72.4 | 0.74(0.32, 1.74) | 0.492 |
| **IPT received** |  |  |  |  |  |  |  |  |  |
| No | 99 | 79.8 | 20.2 | 1.39(0.82, 2.37) | 0.218 | 41.4 | 58.6 | 1.51(0.97, 2.35) | 0.067 |
| Yes | 502 | 73.9 | 26.1 | Ref |  | 31.9 | 68.1 | Ref |  |
| **Deworming medication** |  |  |  |  |  |  |  |  |  |
| No | 515 | 74.0 | 26.0 | Ref |  | 35.5 | 64.5 | Ref |  |
| Yes | 86 | 80.2 | 19.8 | 1.43(0.81, 2.51) | 0.218 | 20.9 | 79.1 | 0.48(0.28, 0.83) | **0.009** |

The relationship between malaria and anemia was linked with both household ownership of an ITN and reported sleeping under an ITN. Women who did not own ITN were more than twice as likely to be moderate/severe anemic (**OR=2.06,** 95%CI:1.18, 3.60), while women who reported not sleeping under an ITN the previous night had almost twice the risk of being moderate/severe anemic (**OR=1.75**, 95%CI:1.14, 2.69). Those women who had not previously delivered were more than twice as likely to be moderate/severe anemic compared to women with one or more births (OR**=2.41**, 95%CI:1.55, 3.77). None of the remaining hypothesized risk factors, such as socioeconomic status, minimum dietary diversity, gestational age, and number of ANC visits, were significantly associated with moderate/severe anemia.

In adjusted logistic regression models that included statistically significant risk factors from bivariate analysis (**Table 3**), tertiary education **(aOR=0.20,** 95%CI:0.04, 0.96) and no previous delivery (**aOR=2.13,** 95%CI:1.18, 3.85) were the only two factors that were significantly associated with anemia (<11.0g/dl) (**Table 3**). The current study found that maternal dietary diversity (<5 MDD-W) was not statistically associated with anemia (aOR=0.80, 95%CI:0.67, 1.38) or moderate/severe anemia (aOR=0.74, 95%CI:0.44, 1.24). However, in using moderate/severe anemia (<9.0g/dl) as the cut-off point for anemia, three explanatory factors were significantly associated. That is, participants whose occupation status was categorized as "other" (e.g. herdsmen) were three times more likely to have moderate/severe anemia (**aOR=2.90,** 95%CI: 1.04, 8.09). Women who had no previous delivery continued to be more likely to have moderate/severe anemia (**aOR=2.13,** 95%CI: 1.28, 3.54). Finally, participants who received deworming medication remained at lower risk of moderate/severe anemia (**aOR=0.51,** 95%CI:0.29, 0.90) than those who did not take these medications.

**Table 3:** Multivariate analysis of anemia status among pregnant women in Northern Ghana (n=601)

|  |  | Anemia<br>(<11.0g/dl) | No anemia<br>(≥11.0g/dl) | Anemia (<11.0g/dl)<br>versus no anemia<br>(≥11.0g/dl) |  | Moderate/severe<br>anemia<br>(<9.0g/dl) | Mild/no<br>anemia<br>(≥9.0g/dl) | Moderate/severe anemia<br>(<9.0g/dl) versus mild/no<br>anemia (≥9.0g/dl) |  |
| --- | --- | --- | --- | --- | --- | --- | --- | --- | --- |
|  | Frequency | % | % | Adjusted OR(CI) | P-value | % | % | Adjusted OR(CI) | P-value |
| **Education status** |  |  |  |  |  |  |  |  |  |
| None | 450 | 76.2 | 23.8 | Ref |  | 30.9 | 69.1 | Ref |  |
| Primary | 46 | 76.1 | 23.9 | 0.96(0.45, 2.03) | 0.908 | 41.3 | 58.7 | 1.48(0.76, 2.88) | 0.253 |
| JSS/Middle School | 53 | 69.8 | 30.2 | 0.59(0.30, 1.18) | 0.135 | 41.5 | 58.5 | 1.31(0.69, 2.49) | 0.407 |
| Senior Secondary | 38 | 73.7 | 26.3 | 0.64(0.27, 1.52) | 0.312 | 44.7 | 55.3 | 1.19(0.54, 2.62) | 0.657 |
| Tertiary | 14 | 50.00 | 50.00 | **0.20**(0.04, 0.96) | **0.044** | 28.6 | 71.4 | 0.75(0.15, 3.71) | 0.723 |
| **Occupation status** |  |  |  |  |  |  |  |  |  |
| Farmer | 233 | 77.7 | 22.3 | Ref |  | 31.3 | 68.7 | Ref |  |
| Petty trader | 181 | 70.2 | 29.8 | 0.72(0.45, 1.17) | 0.185 | 29.8 | 70.2 | 0.95(0.60, 1.51) | 0.840 |
| Salaried worker | 14 | 57.1 | 42.9 | 0.85(0.16, 4.46) | 0.852 | 28.6 | 71.4 | 0.95(0.19, 4.72) | 0.948 |
| Artisan/laborer | 79 | 77.2 | 22.8 | 0.95(0.48, 1.89) | 0.874 | 34.2 | 65.8 | 0.83(0.45, 1.53) | 0.549 |
| No occupation | 75 | 77.3 | 22.7 | 0.93(0.47, 1.86) | 0.844 | 41.3 | 58.7 | 1.32(0.72, 2.41) | 0.367 |
| Others | 19 | 78.9 | 21.1 | 1.17(0.35, 3.87) | 0.797 | 63.2 | 36.8 | **2.90**(1.04, 8.09) | **0.042** |
| **Wealth index** |  |  |  |  |  |  |  |  |  |
| Poorest | 118 | 73.7 | 26.3 | 1.14(0.62, 2.08) | 0.676 | 28.8 | 71.2 | 0.96(0.52, 1.75) | 0.893 |
| Poor | 119 | 76.5 | 23.5 | 1.40(0.77, 2.54) | 0.277 | 34.5 | 65.5 | 1.33(0.74, 2.38) | 0.337 |
| Medium | 120 | 80.0 | 20.0 | 1.80(0.98, 3.31) | 0.059 | 37.5 | 62.5 | 1.46(0.83, 2.57) | 0.191 |
| Wealthy | 122 | 68.0 | 32.0 | Ref |  | 31.2 | 68.8 | Ref |  |
| Wealthiest | 122 | 76.2 | 23.8 | 1.65(0.90, 3.01) | 0.102 | 35.3 | 64.7 | 1.20(0.68, 2.12) | 0.533 |
| **Gestational age (in weeks)** |  |  |  |  |  |  |  |  |  |
| 20-27 | 11 | 72.7 | 27.3 | 1.17(0.28, 4.89) | 0.830 | 36.4 | 63.6 | 1.23(0.34, 4.54) | 0.752 |
| 28-36 | 547 | 75.1 | 24.9 | Ref |  | 33.3 | 66.7 | Ref |  |
| ≥37 | 43 | 72.1 | 27.9 | 0.63(0.28, 1.41) | 0.261 | 34.9 | 65.1 | 1.25(0.58, 2.68) | 0.572 |
| **Dietary diversity** |  |  |  |  |  |  |  |  |  |
| <5 | 91 | 73.6 | 26.4 | 0.80(0.47, 1.38) | 0.427 | 29.7 | 70.3 | 0.74(0.44, 1.24) | 0.249 |
| ≥5 | 510 | 75.1 | 24.9 | Ref |  | 34.1 | 65.9 | Ref |  |
| **Slept under ITN previous night** |  |  |  |  |  |  |  |  |  |
| No | 104 | 77.9 | 22.1 | 1.15(0.67, 1.96) | 0.621 | 44.2 | 55.8 | 1.48(0.93, 2.35) | 0.101 |
| Yes | 497 | 74.3 | 25.7 | Ref |  | 31.2 | 68.8 | Ref |  |
| **No. of previous deliveries** |  |  |  |  |  |  |  |  |  |
| 0 | 142 | 81.7 | 18.3 | **2.13**(1.18, 3.85) | **0.012** | 47.9 | 52.1 | **2.13**(1.28, 3.54) | **0.004** |
| 1 | 122 | 77.1 | 22.9 | 1.57(0.89, 2.75) | 0.116 | 32.8 | 67.2 | 1.21(0.72, 2.03) | 0.477 |
| 2-3 | 214 | 70.1 | 29.9 | Ref |  | 27.6 | 72.4 | Ref |  |
| ≥4 | 123 | 73.2 | 26.8 | 1.10(0.65, 1.84) | 0.728 | 27.6 | 72.4 | 0.97(0.58, 1.64) | 0.916 |
| **Total ANC visits** |  |  |  |  |  |  |  |  |  |
| < 4 | 286 | 76.9 | 23.1 | Ref |  | 33.9 | 66.1 | Ref |  |
| 4-7 | 286 | 72.4 | 27.6 | 0.91(0.61, 1.38) | 0.663 | 33.6 | 66.4 | 1.12(0.76, 1.64) | 0.564 |
| ≥8 | 29 | 79.3 | 20.7 | 1.91(0.64, 5.74) | 0.248 | 27.6 | 72.4 | 0.88(0.32, 2.37) | 0.795 |
| **IPT received** |  |  |  |  |  |  |  |  |  |
| No | 99 | 79.8 | 20.2 | 1.26(0.72, 2.19) | 0.424 | 41.4 | 58.6 | 1.54(0.95, 2.48) | 0.080 |
| Yes | 502 | 73.9 | 26.1 | Ref |  | 31.9 | 68.1 | Ref |  |
| **Deworming medication** |  |  |  |  |  |  |  |  |  |
| No | 515 | 74.0 | 26.0 | Ref |  | 35.5 | 64.5 | Ref |  |
| Yes | 86 | 80.2 | 19.8 | 1.53(0.85, 2.76) | 0.158 | 20.9 | 79.1 | **0.51**(0.29, 0.90) | **0.021** |

## DISCUSSION

We sought to assess the independent contributions of dietary diversity and other predictors to anemia status during pregnancy. Three-quarters of women were anemic (<11.0g/dl), including one-third of whom had moderate/severe anemia. This prevalence of anemia in our study area is considered by WHO to be a severe public health problem ^3^, and is higher than that for Ghana as a whole, and for the Africa region. Indeed, this anemia prevalence is considerably higher than the 44.6% reported in the 2014 Ghana Demographic and Health Survey (GDHS) report. Other studies of pregnant women indicated anemia prevalences of 70.0% in northern Ghana, 62.6% in Southern Ghana and 51.9%-to-59.6% in Africa ^3,7,20,21,22^. Not surprisingly, anemia prevalence among pregnant women varies by geographical area, culture, seasonality and countries ^23^ such as 75% in the current study, compared with 18% and 57% in Ethiopia ^24,25^. The high prevalence that we observed might be due to increased physiological demands for nutrients during the second and third trimesters of pregnancy ^26^. It could also be attributed to increased demand for iron by the growing fetus, particularly during the last trimester of pregnancy ^27^. In addition to these possible explanations, the specific causes underlying the high anemia prevalence in the study setting might be attributed to the geographical area and cultural preferences, as was seen in northern Ghana with prevalences of 47% mild, 20% moderate, and 3% severe ^10^.

In bivariate analysis, some socio-demographic factors (e.g. older age) were associated with lower odds of moderate/severe anemia among pregnant women, while others such as occupation (e.g. herdsmen) and parity (primigravidae) were associated with higher odds of moderate/severe anemia. Our finding that increased age and greater parity were associated with lower risk of moderate/severe anemia may be due to the experience acquired by women during previous pregnancies and with ANC services, for example in the use of ITN and IPT. A similar study conducted in northern Ghana also found that increased parity was associated with reduced anemia, – ranging from 75.0% anemia among primigravidae to 43.7% among multigravidae ^20^. Also, another investigation in southern Ghana reported mean Hb concentrations of 9.7g/dl for primigravidae, 10.1g/dl for secundigravidae and 10.5g/dl for multigravidae ^28^. However, other studies have found that higher parity (multigravidae) was associated with greater risk of anemia ^25,29,30,31^. This might be due to differences in birth spacing rather than total parity. For example, pregnant women with short pregnancy intervals were more likely to develop anemia during pregnancy than those with longer birth spacing ^27^.

Our results indicated that pregnant women who worked in other occupations (e.g. herdsmen) were are at greater risk of moderate/severe anemia. Usually, these families live in the peripheral settings of rural Ghana communities, and therefore have reduced access to health facilities. In addition, they are often from minority ethnic groups, hence are more likely to delay ANC initiation until second or third trimester of pregnancy. Thus, they fail to exploit all opportunities offered by the ANC ^32^. Salaried workers and traders had lower odds of moderate/severe anemia when compared to farmers. This may be due to their increased access to food, greater dietary diversity and improved food security compared to farmers, for example, whose livelihood depends on seasonal crops to meet their nutrient requirements.

Greater parity was associated with lower odds of moderate/severe anemia in multivariable analyses (Table 3). Specifically, first-time pregnant women were more than twice as likely to be moderate/severe anemic. This is consistent with previous studies where primigravidae had increased risk of anemia compared to multigravidae ^12^. Until uncertain rituals are performed by the family members of the husband to publicly declare or announce the pregnancy, they are unwilling to start ANC visits for fear of miscarriage or other complications.

Dietary diversity may be an indicator of food access and use, and of diet quality. We observed that after controlling for potential confounding factors (maternal education, occupation, wealth index, ITN ownership and utilization, parity, ANC attendance, IPT and deworming medication) in multivariate logistic regression model, achieving the MDD-W indicator was not associated with moderate/severe anemia. Previous research in Ghana found consumption of other fruits or and seeds/nuts was statistically associated with anemia and MDD-W was not associated with anemia ^10^, but another study found that high maternal MDD-W was associated with a decreased risk of anemia ^20^. Diet is an important factor for anemia. Some eating patterns or habits may predispose pregnant women to an increased risk of developing anemia. Poor dietary diversity leads to inadequacies in minerals and vitamins. Though we observed that 85% of the pregnant women consumed ≥5 food groups in the previous day, dietary diversity was not associated with a higher Hb concentration. This lack of association between MDD-W and moderate/severe anemia may be due to inappropriate kinds of foods (i.e. animal-source foods with highly bioavailable iron) that would be linked to lower anemia risk or challenges of nutrients bioavailability from the consumed diets. We also found that consumption of animal-source foods was not associated with lower odds of anemia in the current study.

Overall, 91% of pregnant women reported of ITN ownership and 83% slept under the ITN the night before the interview. The coverage and utilization of ITN higher than a previous study conducted in southern Ghana where 75% of pregnant women reported ownership of ITN and only 49% slept under it the night preceding enrolment ^33^. Similarly, the coverage of at least one IPT dose (84%) in the current study exceeds the national average (68%) ^33^. Independent of malaria interventions (household ownership of ITN and slept under ITN on previous night), ≥4 ANC visits and deworming medication were associated with lower odds of moderate/severe anemia among pregnant women aged ≥20 weeks of gestation. In our study, deworming medication was protective against moderate/severe anemia. This varies from the a previous study which reported that iron supplementation and deworming treatment was not significant associated with anemia ^30^. We observed that pregnant women who received IPT (83%) had lesser of risk to develop moderate/severe anemia. An explanatory factor such as IPT was not associated with moderate/severe anemia. Our current finding is similar to a study conducted in Ghana which found that IPT shown a weak association with anemia using bivariate analysis but no association when adjusted for confounders in multivariate analysis ^20^. In a previous study conducted in Ethiopia, malaria infection during pregnancy had higher risk to develop anemia ^27^. Our study found that ownership of ITN and utilization of ITN were associated with reduced odds of moderate/severe anemia. This is suggestive of effective intervention in the study area. Deworming medication was associated with moderate/severe anemia. Parasitic infections are known to be a major cause of anemia. Thus, treatment against intestinal parasites help to improve hemoglobin concentration ^3^.

### Strengths and Limitations

In terms of strength, the study focused on an important category of pregnant women with implications on birth outcomes. The sample size was also large and representative of the population accessing ANC services at HFs. This is probably the first study to assess the predicators of moderate/severe anemia among pregnant women (≥20 gestational weeks) in northern Ghana.

This study had several limitations and the findings should be interpreted as such. The study design was cross-sectional and does not allow us to assess causality. In addition, participant recall bias was a limitation but the interviews were conducted using a structured questionnaire and the interviewers were trained nurses and health science educators to mitigate the effect of recall bias. The study sample was limited to pregnant women who received ANC services at HFs, thus, some pregnant women were excluded. This may not be reflective of the study setting. However, the majority of Ghanaian women now seek ANC services.

## CONCLUSION

The prevalence of anemia among pregnant women was nearly twice as national estimate in Ghana, and varied by maternal age, parity and occupation. Minimum dietary diversity was not associated to moderate/severe anemia. Household ownership of ITN, use of ITN and deworming medication were associated with moderate/severe anemia. There is the need to intensify education on ANC services, ITN utilization and deworming medication to prevent and reduce anemia prevalence among pregnant women.

## Abbreviations

ANC: Antenatal Care
aOR: Adjusted Odds Ratio
CI: Confidence Interval
DHS: Demographic and Health Survey
FAO: Food and Agricultural Organization
GDP: Gross Domestic Product
GHS: Ghana Health Service
GHS: Ghana Cedis
GHSERC: Ghana Health Service Ethics Review Committee Hb Hemoglobin
HC: Health Centre
HF: Health Facility
HIV: Human Immuno-Deficiency Virus
IPT: Intermittent Preventive Treatment
ITN: Insecticide Treated Mosquito Net
MDD-W: Minimum Dietary Diversity for Women OR Odds Ratio
PCA: Principal Component Analysis
RCH: Reproductive and Child Health
TB: Tuberculosis
USD: United State Dollars

## Acknowledgements

We are grateful to the two research assistants and staff of Ghana Health Service who participated in the data collection. In addition, we acknowledge financial support from PARTNER II through funding from the Fogarty International Center of US National Institutes of Health (D43TW009353).

## Data Availability Statement

The dataset used and/or analyzed during the study is available from the corresponding author upon request.

## Author contributions

MNA and MW conceptualized and designed the protocol and received support from AJ and RA. MW and RA supervised the implementation of the study. MNA and MY conducted the study. MNA and MW drafted the manuscript. RA, AJ and MY edited the manuscript. All authors read and approved the final manuscript.

